# Modular analysis of a logical regulatory network reveals how Fe-S cluster biogenesis is controlled in the face of stress

**DOI:** 10.1101/2022.07.15.500164

**Authors:** Firas Hammami, Laurent Tichit, Frédéric Barras, Pierre Mandin, Elisabeth Remy

## Abstract

Iron-sulfur (Fe-S) clusters are important cofactors conserved in all domains of life, yet their synthesis and stability is compromised in stressful conditions such as iron deprivation and oxidative stress. Two conserved machineries, Isc and Suf, assemble and transfer Fe-S clusters to client proteins. The model bacterium *Escherichia coli*, where Fe-S biogenesis has been extensively studied, possesses both Isc and Suf machineries and their utilisation is under the control of a complex regulatory network. To better understand the underlying regulatory mechanisms and the dynamics behind Fe-S biogenesis in function of environmental conditions, we built a logical model describing Fe-S biogenesis in *E. coli*. This logical model is centered on three modules : 1) the Fe-S biogenesis module containing the Fe-S cluster assembly machineries Isc and Suf and the transcription factor IscR, the main regulator of Fe-S homeostasis; 2) the iron homeostasis module containing the free intracellular iron regulated by the iron sensing regulator Fur and the non-coding regulatory RNA RyhB involved in iron sparing; 3) the oxidative stress module representing intracellular H_2_O_2_ accumulation, which activates OxyR, the regulator of catalases and peroxidases that decompose H_2_O_2_ and limit Fenton reaction. Inputs of the model represent extracellular iron and oxygen environmental conditions, whereas ErpA, NfuA (Fe-S carrier proteins), and Suf are output nodes of the model. Analysis of this comprehensive model reveals 5 different types of system behaviours depending on environmental conditions, and provides a better understanding on how oxidative stress and iron homeostasis combine and control Fe-S biogenesis. Using the model, we were able to predict that an *iscR* mutant would present growth defects in iron starvation, due to partial inability to build Fe-S clusters, and we validated this prediction experimentally.

**Author summary:** Iron sulfur (Fe-S) clusters appeared at the origins of life, when oxygen tension was low and iron plentiful, and have been used since as important cofactors for a wide variety of proteins involved in a plethora of reactions. However, synthesis and stability of Fe-S clusters is compromised in conditions where iron is low or in presence of reactive oxygen species. Living organisms have developed complex regulatory network to allow biogenesis of Fe-S clusters in function of environmental conditions. Thus, understanding this regulation is of primary importance as changes in Fe-S cluster biogenesis impact the physiology of the cell and is for instance involved in resistance of bacteria to certain antibiotics. To do this, we used a modeling approach to gain a global systemic understanding of the process. We thus developed a mathematical logical model which extensively describes the regulatory network that controls biogenesis of Fe-S clusters in the model bacterium *Escherichia coli*. Analysis of the model reveals how Fe-S biogenesis is organized in function of environmental conditions and reveals how oxidative stress and iron homeostasis combine and control Fe-S biogenesis.

## Introduction

Iron sulfur clusters (Fe-S clusters) are ubiquitous prosthetic groups formed of two to four iron centers covalently bound by sulfur atoms. Fe-S clusters containing proteins are present in virtually all living organisms and are involved in a plethora of biological roles ranging from respiration to the TCA cycle, gene regulation or bacterial virulence [1]. However, Fe-S clusters are extremely sensitive to environmental stresses, in particular iron deprivation and oxidative stress. Indeed, iron is needed to assemble Fe-S clusters, while oxidative stress generates Reactive Oxygen Species (ROS) that destroy Fe-S clusters and inactivate Fe-S proteins, such as dehydratases [2–4]. Moreover iron and ROS are linked through the Fenton reaction: Fe^2+^ + H_2_O_2_ → Fe^3+^ + HO^−^ + HO_·_. This reaction leads to the production of hydroxyl radicals which cause DNA damage and presents no known direct scavenger.

Fe-S cluster biogenesis is thus tightly regulated by actors of the iron homeostasis and oxidative stress responses in a complex regulatory network, involving multiple regulators of different nature (transcription factors and regulatory RNAs), which makes it difficult to predict how the cell will adapt its Fe-S clusters biogenesis in function of the multiple stresses encountered in nature.

Control of Fe-S clusters biogenesis has been best studied in the model bacterium *Escherichia coli* which can adapt to multiple niches wherein iron and oxygen concentrations can vary greatly. In this bacterium, Fe-S clusters biogenesis is ensured by two main machineries, Isc and Suf, present in all domains of life [1, 4]. Here we aimed at deciphering the regulatory network controlling Fe-S clusters biogenesis in *E. coli* through a mathematical modelling approach. Such models reveal the global dynamical behaviour emerging from the network and their analysis untangles the contribution of the different regulators, identifies key actors, and allows to make predictions, e.g., on how biological systems respond to environmental cues or to mutations in genes of the system.

While the Isc and Suf machineries consist of different macromolecular complexes, their general mechanism is similar. In the first step, iron and sulfur are acquired (through a yet unknown iron donor, and from cysteine, respectively) and then assembled on a scaffold complex. The preformed Fe-S cluster is then delivered to apo-proteins through A-Type carriers (ATC) [1]. In addition to ATCs of the main Fe-S biogenesis machineries, IscA and SufA, *E. coli* displays non-Isc non-Suf carriers such as ErpA and NfuA that serve to multiply the Fe-S cluster transport routes in function of environmental conditions [5]. In particular, *erpA* is essential in aerobiosis, while the *nfuA* mutant is extremely sensitive to iron starvation and oxidative stress [6, 7].

In *E*.*coli* Fe-S biogenesis is mainly regulated by the IscR transcription factor, itself an Fe-S cluster containing protein matured almost exclusively by the Isc machinery [8–12]. In its Holo (Fe-S bound) form IscR represses the transcription of several genes among which the *iscRSUA* operon, encoding the Isc machinery, *erpA* and *nfuA*. In contrast, accumulation of apo-IscR form activates transcription of the *sufABCDSE* operon, encoding the Suf machinery. In this way, both the Fe-S cluster state and the concentration of IscR dictate the choice of Fe-S biosynthesis machinery and carriers.

Other global regulators are involved in this choice, linking iron homeostasis and oxidative stress to Fe-S cluster biogenesis. In *E. coli*, iron homeostasis is mainly controlled by the Fur transcription factor and by the non-coding regulatory RNA RyhB [13]. When bound to iron, Fur represses iron import genes and inhibits Suf and RyhB expression [14]. Iron starvation alleviates Fur mediated repression of the aforementioned genes. RyhB induction causes iron sparing, both by inhibiting translation of non-essential Fe using proteins and by promoting siderophores production [13–16]. RyhB basepairs to the *iscRSUA* mRNA and inhibits expression of the Isc machinery while leaving that of IscR intact. In this way RyhB was proposed to indirectly lead to the accumulation of apo-IscR and to Suf expression.

Among the several oxidative challenges that can affect Fe-S clusters, we focus on H_2_O_2_ since it can react with the Fe of the Fe-S cluster and it is the ROS involved in Fenton reaction. Response to H_2_O_2_ driven oxidative stress is mainly regulated by OxyR in *E. coli*. OxyR activation by H_2_O_2_ leads to the induction of the Suf machinery. In addition, OxyR also activates the expression of the catalase KatG and peroxidase AhpCF (together annotated Hpx), that diminish H_2_O_2_ concentration, as well as the expression of Dps and YaaA, that store the free intracellular iron in order to limit Fenton reaction [10,17,18]. Moreover, OxyR induces Fur expression to limit the ROS-dependent demetallation of Fur [19].

Mathematical modelling is increasingly used to formalize, synthesize and analyse set of regulatory interactions controlling biological processes. We present here a qualitative mathematical model encompassing the molecular actors of Fe-S biogenesis regulation, iron homeostasis and oxidative stress response based on the logical formalism. This model explains mechanistically how the two main systems Isc and Suf are coordinated in response to external iron and oxygen signals. We then determined the individual and combined contributions of oxygen and iron on Fe-S cluster biogenesis, before predicting situations of Fe-S dependent growth defects in mutants.

## Results

### Establishment of a logical model for the Fe-S cluster biogenesis

We built a gene regulatory network describing Fe-S cluster biogenesis regulation in response to iron and oxygen concentrations in the environment. It consist in an directed signed graph where nodes represent genes or biological components, and edges the positives or negative influences (see Materials and Methods). The model encompasses the three biological processes described above: iron homeostasis, with nodes *Fe*_*free*_, *Fur* and *RyhB* components; oxydative stress with nodes *OxyR,Hpx* (which represents the catalases and peroxidases involved in H_2_O_2_ detoxification) and node *H*_*2*_ *O*_*2*_; and Fe-S clusters biogenesis with the nodes *IscR-A, IscR-H, Isc, NfuA, ErpA* and *Suf*. Input nodes *Fe*_*ext*_ and *O*_2_ stand for external iron and oxygen. Note that we chose to describe the IscR regulator with two nodes, *IscR-A* (for apo IscR) and *IscR-H* (holo-IscR) to describe both the expression and the maturation of the protein. The three output nodes *ErpA, NfuA* and *Suf*, and also node *Isc* serve as read-out of Fe-S biogenesis. The resulting model is composed of 14 nodes and 28 directed interactions (see Fig.1). We have focused on the three topological modules that correspond to the three biological processes included in the model.

**Fig 1.**
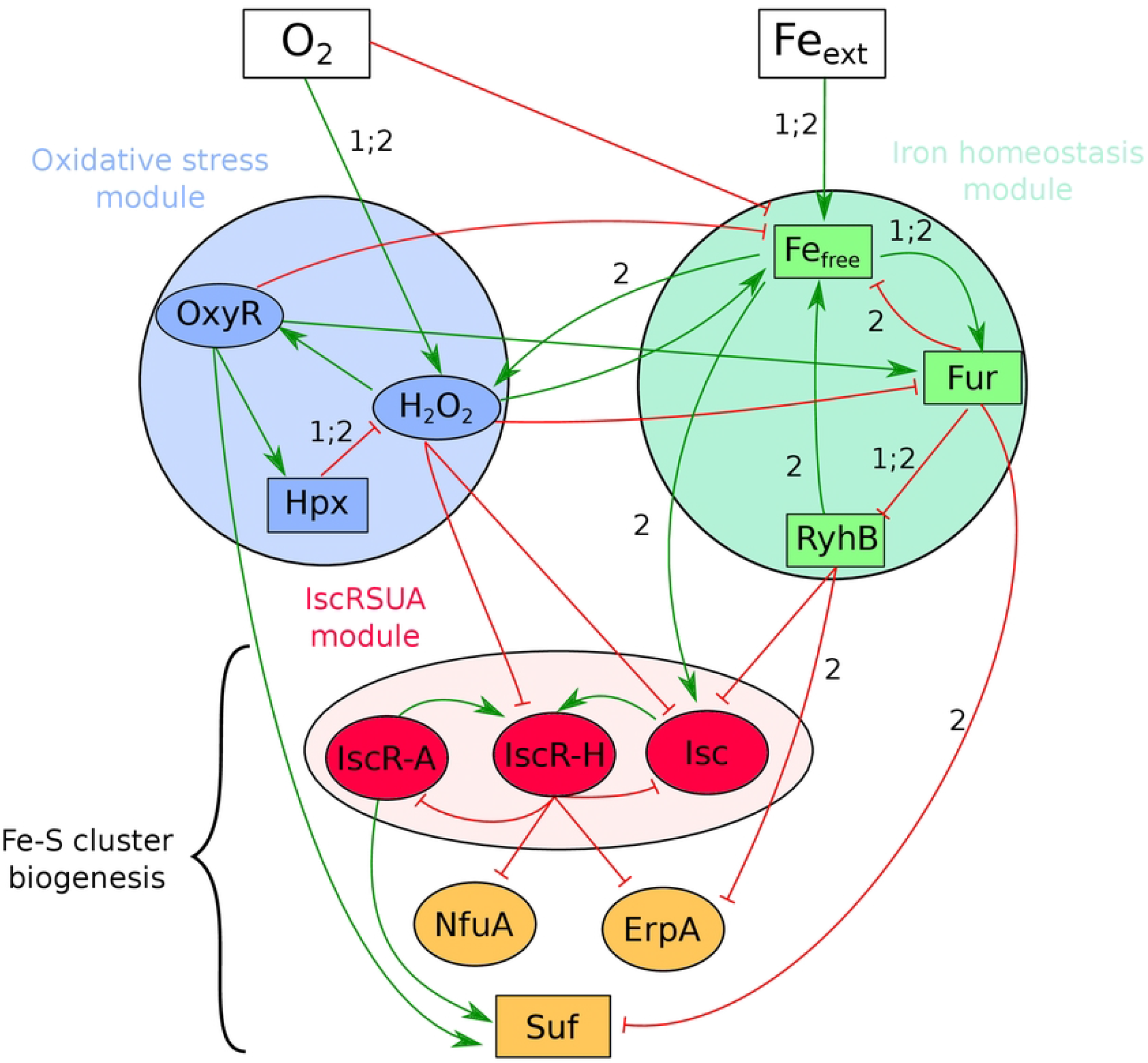
Regulatory Graph centered on the Fe-S biogenesis in *E. coli*. Nodes represent biological components, and edges regulations: activations (green, normal arrow) or inhibitions (red, T-shaped arrows). Ellipses stand for Boolean nodes, rectangles multivalued nodes. Labels on edges specify the regulatory thresholds (equal to 1 if not indicated); two labels on the same edge means the regulation happens at both thresholds. Each module is represented by a color: red nodes belong to module Isc; blue nodes to the Oxidative stress response module; green nodes to the Iron homeostasis module. The three orange nodes are the outputs of the model. The shadowed ellipse gathers nodes of the biological Fe-S cluster biogenesis module (Isc module and outputs).

We parameterized this graph to integrate its dynamics (see Materials and Methods). Half of the components are associated with Boolean variables (representing the inactivity/activity state of the components), and half with ternary variables representing low, medium and high activity levels, except for two nodes: the *O*_2_ node for which the 3 levels indicate aerobic, anaerobic and oxidative stress conditions; the *Hpx* node for which the basal level is 1 (representing constitutive expression of catalases that scavenge low doses of H_2_O_2_ produced by respiration), level 2 represent high expression of catalases during oxidative stress, and level 0 is used to simulate Knocked Out mutation. Logical equations are settled on the basis of available biological knowledge (see Table 1, S1 Text and S1 Table). For example, the node *IscR-H* is activated by the presence of both *IscR-A* and *Isc* in the absence of *H*_*2*_ *O*_*2*_ node. The annotation of each node and edge, associated with bibliographical references and documentation, are included in the GINsim model file (cf Materials and Methods and the ginml file). The attractors of the model are described in S2 Table. The model was validated by comparing predictions of the model to literature data that was not used for model construction (see S3 Table).

**Table 1.**
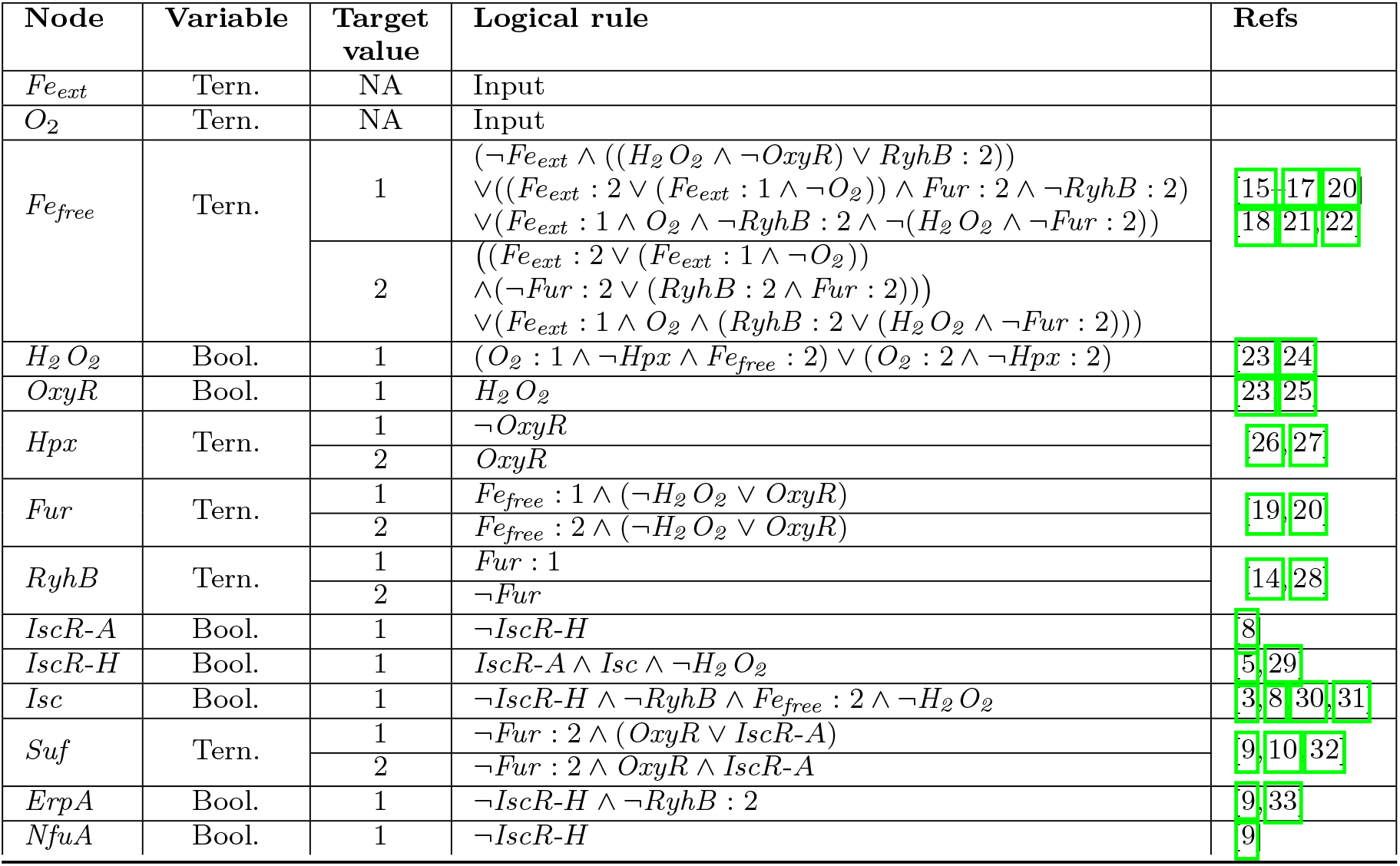
Logical rules associated with each node of the model (except the input nodes, for which the initial values are fixed during the simulations). In column 2 is precised if the associated variable is ternary or Boolean. For each level (or target value, column 3) is given the logical rule to be satisfied (column 4). The literature references are displayed in column 5.

### The model reveals five qualitative asymptotical behaviours depending on environmental constraints

We performed simulations of the model and characterized the attractors in order to capture the asymptotical behaviours of the system for each of the nine environmental conditions (see Materials and Methods). Each combination of input values determines a unique attractor, which is a stable state when both input nodes values *Fe*_*ext*_ and *O*_2_ are fixed to 1, and a cyclic attractor for each of the 8 others combinations of input values. A complete description of the attractors is provided in S2 Table. A global visualisation of the asymptotic activities generated by the model (in terms of oscillations or stability) at the module scale highlights clearly how the regulatory processes evolve and adapt to the environment (cf Fig 2, A). These modules form well-known topological motifs (a circuit for the Oxidative stress module [34], a chorded circuit for the iron homeostasis [35] and a 2 petals-flower for the IscRSUA module [36]), that are 3-components motifs constituted of negative feedback loops known to potentially control oscillations [37]. Note that a theoretical study of these motifs help to explain the global behaviour of the system [38].

**Fig 2.**
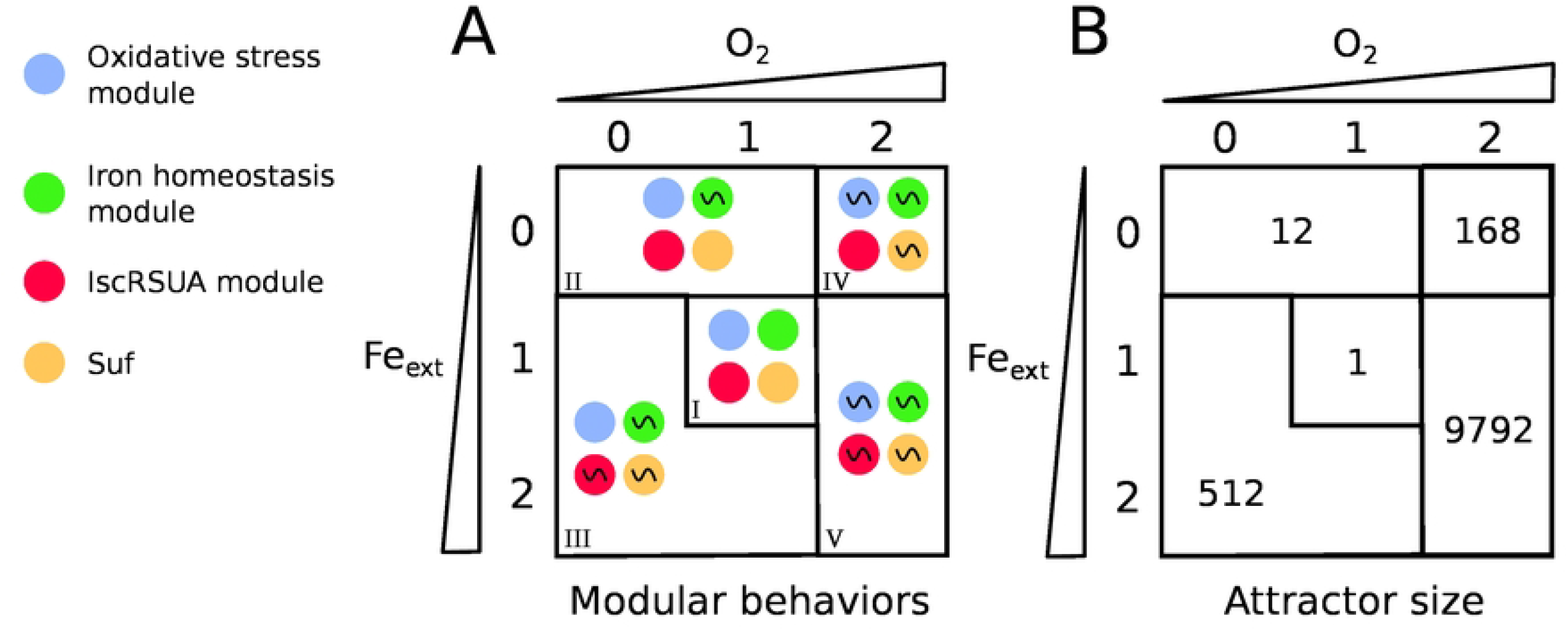
Modular description of the five asymptotic classes of the model. The two axes of the square represent the environmental conditions *O*_2_ and *Fe*_*ext*_ taken their values between 0 and 2. The cutting of the grid in areas separates the five different classes of asymptotical behaviours of the system. **A**: Each area contains four circles representing the behaviour of the four modules of the RG identified by a color: Oxydative stress response (blue), iron homeostasis module (green), Isc module (red), Suf (yellow). The symbol ~ denotes an oscillatory asymptotical behaviour of the module, otherwise it is stable. **B**: The number *m* indicates the size of the cyclical attractor (number of states).

Five classes of behaviours emerge (numbered from I to V in Fig. 2, A), characterized by the oscillating modules: the system is stable when the inputs are maintained to their intermediate level (class I); only the Iron module oscillates in case of external iron starvation with no oxidative stress (class II); all modules oscillate except the oxidative stress in case of presence of iron and no oxydative stress (class III); all modules oscillate except IscRSUA when iron is absent and oxydative stress present (class IV); all modules oscillate in presence of both iron and oxydative stress (V). The size of the attractors - i.e. the number of states they contain-correlates with the number of oscillating modules and the amplitude of the oscillations in case of multilevel variables (Fig. 2, B). The size of the attractors reflects the number of states involved in the equilibrium of the system for a given input. Somewhat counter-intuitively, the situation combining both iron starvation and oxidative stress (IV) does not display the largest attractor size. In fact, the largest attractor size appears in presence of iron and oxidative stress (V).

### Modular analysis of the model highlights the global adaptation of the system to environmental constraints

We leveraged the specific modular topology of the regulatory graph to analyse the adaptation to the environment. The three biological processes included in the model are represented by small strongly connected modules. Nodes *Fe*_*free*_, *H*_*2*_ *O*_*2*_ and *Isc* have been chosen as representatives of the iron homeostasis, oxidative stress and IscRSUA modules respectively. We also considered nodes *Suf, NfuA* and *ErpA*, outputs of the model, to fully describe Fe-S clusters homeostasis. The asymptotic behaviour of each node is displayed as a heatmap (Fig 3). This format makes it easy to visualise the differences in the behaviour of each process under different environmental conditions.

**Fig 3.**
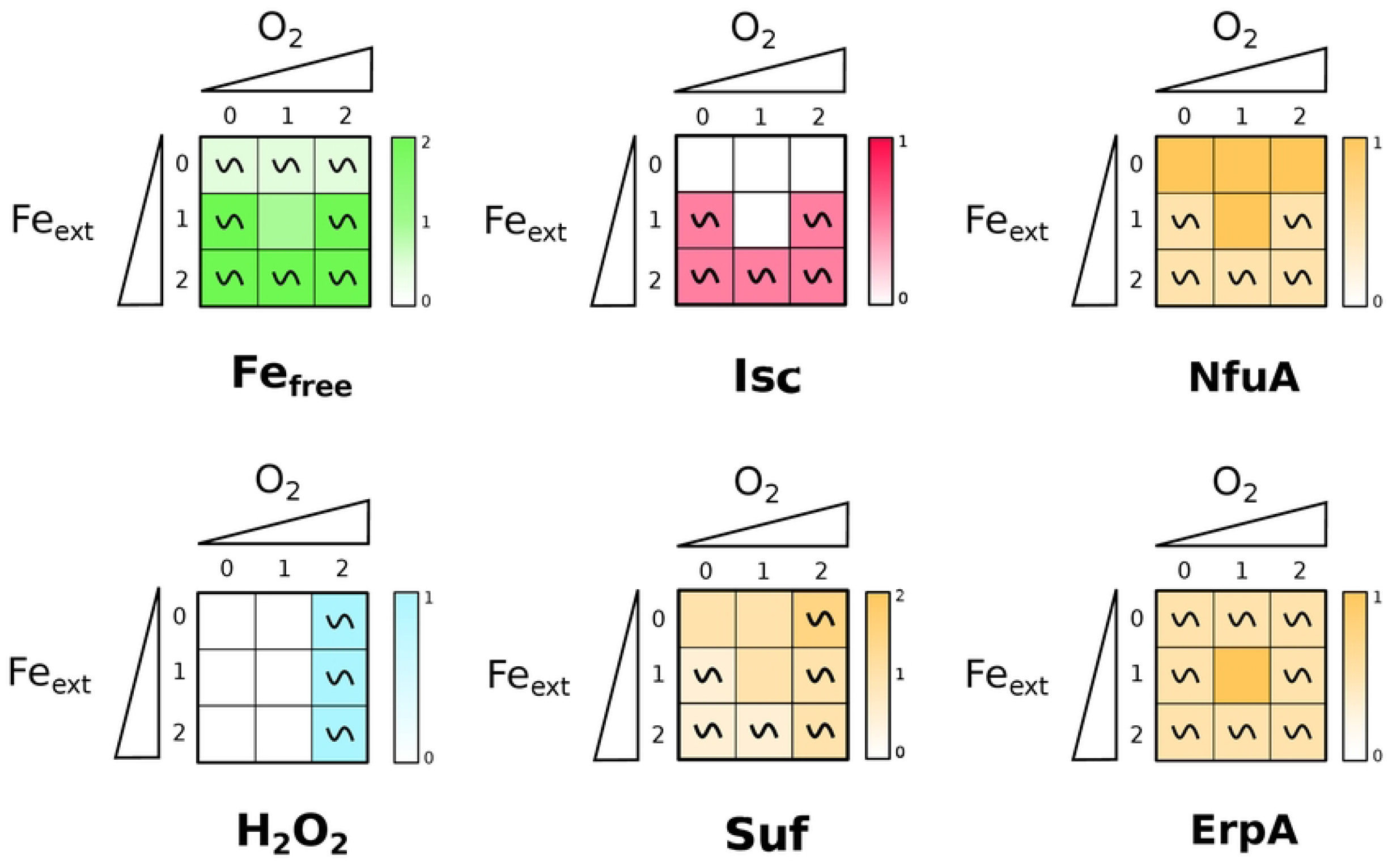
WT simulations. The results of simulations for the six representative nodes in each of the 9 input combinations are shown in the form of a heatmap. Each cell of the grid stands for one of the nine input conditions (depending on the values of *O*_*2*_ and *Fe*_*ext*_). The qualitative asymptotical behaviour of the node is indicated: oscillation (~) or stable (no indication). Its expression level is reflected through the color graduation. In cases of oscillations, the mean over all the states of the cyclic attractor is indicated.

The *Fe*_*free*_ node is stabilized at its medium level 1 when *O*_*2*_ =1 and *Fe*_*ext*_ =1, and presents oscillations in the other environmental conditions, tending to the value 1 (increasing between 0 and 1 in iron starvation; decreasing between 2 and 1 otherwise). This reflects the need for the cell to keep its iron concentration finely tuned around an optimal concentration (*Fe*_*free*_ =1, which represents 10 *μ*M) that allows Fe usage for biological functions while avoiding Fe toxic accumulation in the cell.

The *H*_*2*_ *O*_*2*_ node oscillates in oxidative stress condition (*O*_*2*_ =2, that corresponds to presence of more than 2 *μ*M of H_2_O_2_ in the environment) regardless of the *Fe*_*free*_ value, and is otherwise at 0 in all other situations. This behavior was expected since low doses of H_2_O_2_ are easily scavenged by catalases and peroxydases represented in the model by the Hpx node.

As expected, *Isc* and *Suf* nodes display very distinct behaviors in the WT strain. The *Isc* is active only in the presence of iron (and independently of O_2_), which is in agreement with biochemical data that have shown the Isc machinery to be sensitive to iron chelation [39]. *Suf* node is always present albeit at low levels (oscillating in between 0 and 1) in unstressed conditions corresponding to *Fe*_*ext*_ =1 and *O*_*2*_ =1. The model predicts that its activity increases with increasing *O*_*2*_ and decreasing *Fe*_*ext*_ concentrations. It reaches its maximum level in case of both iron starvation and oxidative stress. Interestingly, it should be noted that this situation is not the one with the largest attractor size (see Fig 2).

*ErpA* and *NfuA* nodes were also predicted to have distinct behaviors, which is mainly due to the fact that *RyhB* node is a regulator of *ErpA* node, but not of *NfuA* node. *NfuA* node presents a complementary behaviour to *Isc* node, with *Isc* being off when *NfuA* is at its maximal values. The prediction that NfuA is more expressed when iron is limiting is in agreement with the essential role that displays this carrier in such conditions [7, 40]. The *ErpA* node displays oscillations in all conditions except for the *Fe*=1 and *O*_*2*_ =1 situation where it is set to its maximum value 1. Both carriers are predicted to be present in line with their role at interacting with both machineries and delivering Fe-S clusters to apo-proteins in most conditions [6, 7, 40].

### Perturbations of the oxidative stress module affect response of the iron module but not conversely

To gain a better understanding of how the oxidative stress and the iron homeostasis modules interact and influence Fe-S biogenesis, we simulated simple or combination of Knocked Out perturbations (KO) affecting one or several modules. Such single or multiple mutants can be easily defined with the software GINsim (see Materials and Methods).

We ran KO perturbation of *OxyR* node (affecting Oxydative stress module), of *Fur* node (iron homeostasis module), and of both. Fig 4 shows the asymptotical behaviours of *H*_*2*_ *O*_*2*_ and *Fe*_*free*_ nodes in WT in these three perturbated situations. Expectedly, mutations affect the module to which the node belongs by blocking the oscillations. Nevertheless, it is interesting to note that the *OxyR* KO simulation, in addition to show an expected increase in *H*_*2*_ *O*_*2*_ (its production can no longer be counteracted), also displays high values of *Fe*_*free*_ node. This behaviour is explained by H_2_O_2_ attack of iron containing proteins and inactivation of Fur. In contrast, while *Fur* KO simulations show an increase of *Fe*_*free*_ in all conditions, reaching its maximal as soon as *Fe*_*ext*_ 1, *H*_*2*_ *O*_*2*_ seems not to be affected by this perturbation. This is due to presence of catalases and peroxidase represented by the *Hpx* node that limit H_2_O_2_ accumulation. We note however that, with such high levels of *Fe*_*free*_, even the transient production of H_2_O_2_ in the *Fur* KO mutant (see *O*_*2*_ =1 and *Fe*_*ext*_ ≥ 1 in Fig.4) will result in production of toxic ROS via the Fenton reaction that account for the growth defect of *fur* mutants.

**Fig 4.**
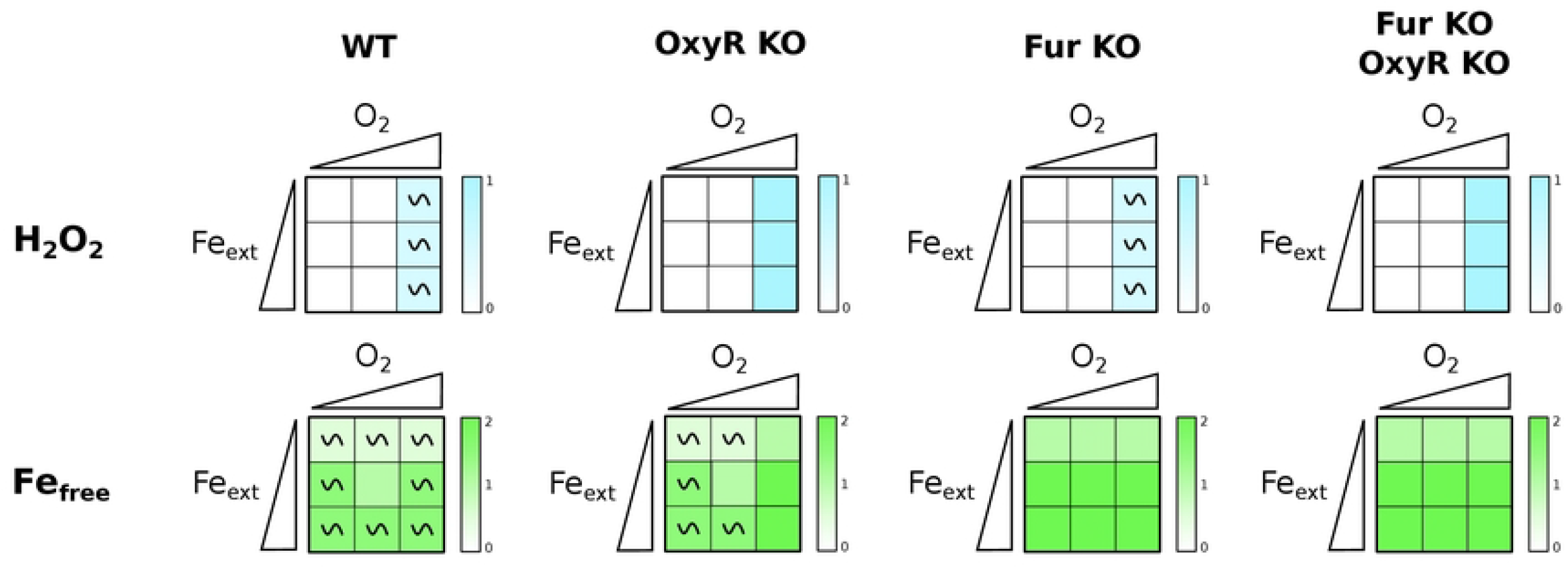
Simulations of simple and double perturbations of iron and H_2_O_2_. Heatmap representation of the asymptotical behaviour of nodes *H*_*2*_ *O*_*2*_ (top line) and *Fe*_free_ (bottom line) in the nine input conditions (see caption of Fig 3) for, from left to right, WT, OxyR KO mutant, Fur KO mutant and Fur KO-OxyR KO double mutant.

Finally, the double perturbation is a combination of the two behaviours of the single mutants, ie the H_2_O_2_ displayed the same behaviour as in the *OxyR* KO single mutant and the *Fe*_*free*_ node behaviour was identical to that of the *Fur* KO mutant. This is not surprising as in this double mutant the homeostatic regulators of each module are deleted. In this double mutant accumulation of H_2_O_2_ and free iron should result in growth defects of this strain due to Fenton reaction seen in known biological situations.

### Perturbations in Fe homeostasis and oxidative stress induce Suf

We next examined the effect of perturbing the oxidative and/or the iron homeostasis module on Fe-S biogenesis by looking at the *Isc* and *Suf* nodes (Fig 5). Simulations with the *OxyR* KO mutant show that this mutation affects both *Isc* and *Suf* behaviours when *O*_*2*_ =2. Indeed, *Isc* expression is predicted to be turned off in this situation (owing to continuous ROS production, see Fig 4), leaving only the *Suf* node present, albeit at diminished levels as compared to the WT situation (*Suf*=1 for any *Fe*_*free*_ value when *O*_*2*_ =2). Strikingly, simulations with *Fur* KO mutant show that *Isc* activity is completely turned down in all situations (due to the constant *RyhB* repression on the *Isc* node), while *Suf* expression is predicted to be turned up in this case. Perturbing both modules retained these features with *Isc* activity turned off (*Isc*=0) and Suf stably turned on (*Suf*=1) in all situations. In conclusion, perturbation of either module decreases activity of *Isc* leaving *Suf* as the functional Fe-S biogenesis system, in agreement with the accepted role of Suf as the only stress response system.

**Fig 5.**
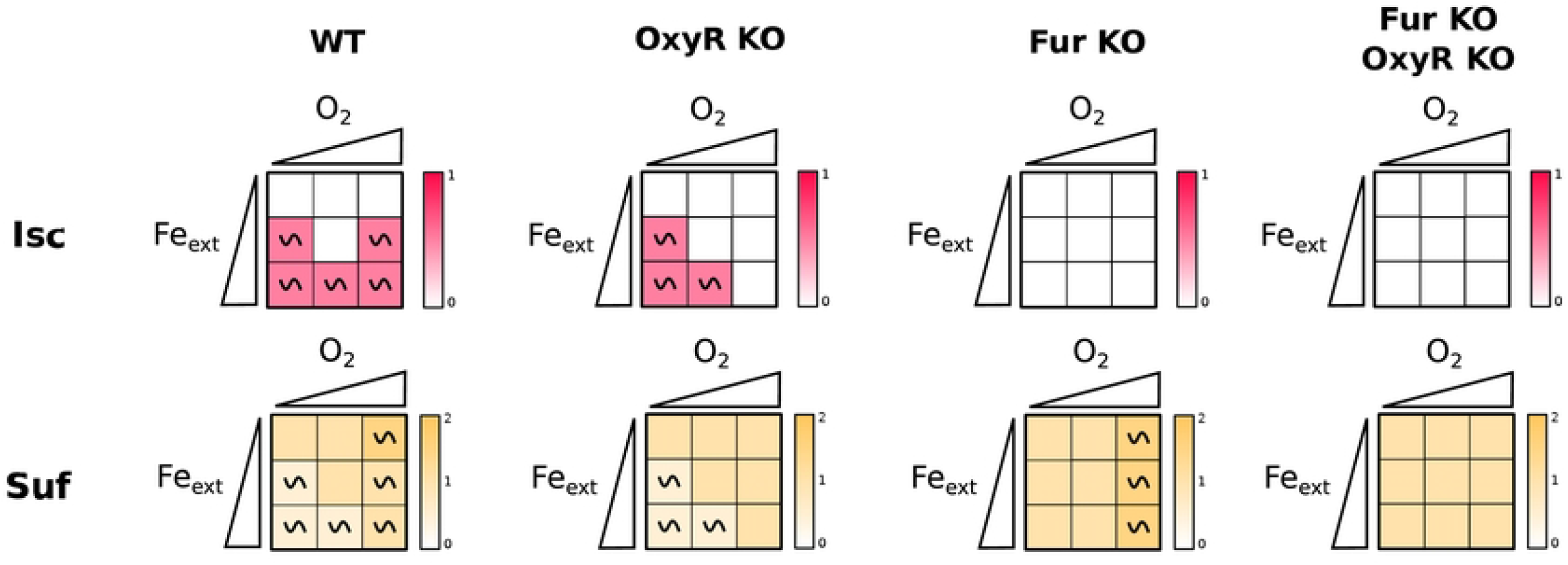
Modules perturbations effects on the Fe-S cluster biogenesis machineries. Heatmap representation of the asymptotical behaviour of nodes *Isc* (top line) and *Suf* (bottom line) in the nine input conditions (see caption of Fig 3) for, from left to right, WT, *OxyR* KO mutant, *Fur* KO mutant and *Fur* KO-*OxyR* KO double mutant.

*ErpA* and *NfuA* nodes, representing the ATCs, were diametrically affected in the *OxyR* and *Fur* KO simulations (Fig 6). The *ErpA* node was turned off (*ErpA*=0) when *O*_2_=2 in the *OxyR* KO mutant, and even more strikingly it was predicted to be off in all conditions in the *Fur* KO mutant. In sharp contrast, the *NfuA* node was constitutively on (*NfuA*=1) in the same conditions. Thus, the model further indicates that NfuA is mainly a stress response ATC as opposed to ErpA which expression is predicted to be turned off in response to stress perturbations.

**Fig 6.**
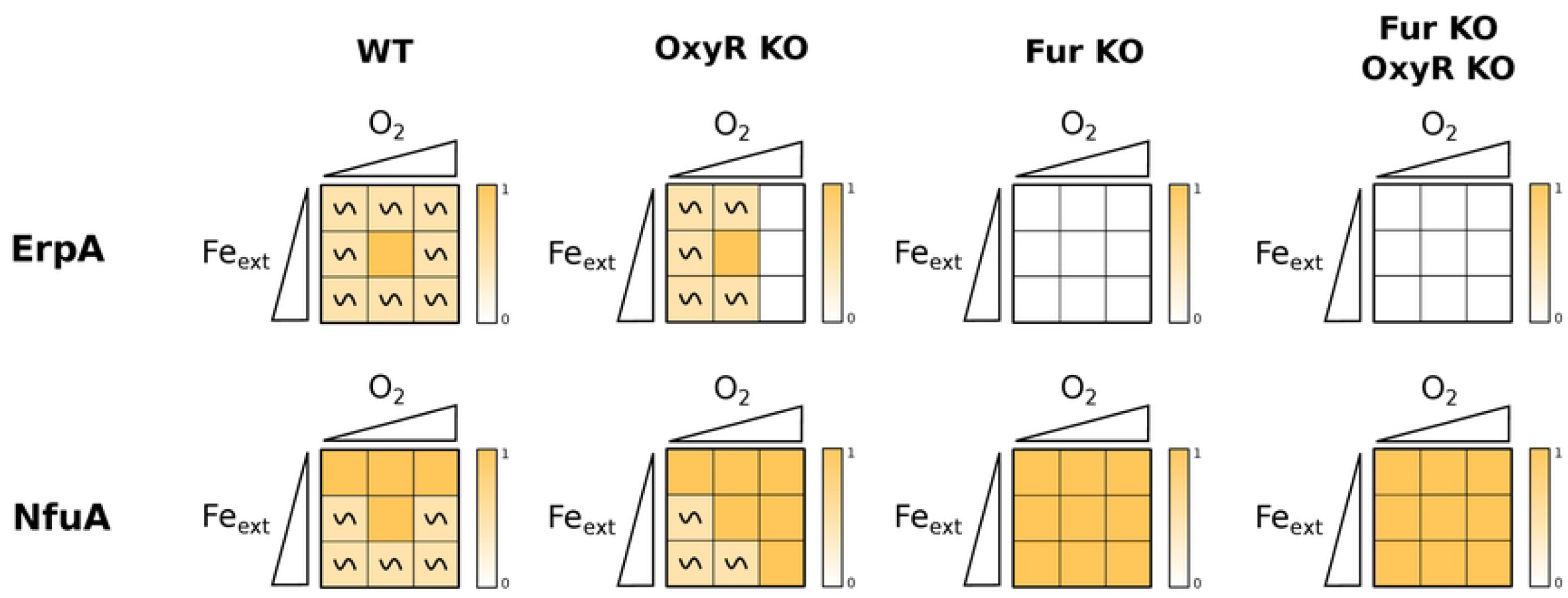
Modules perturbations effects on the ATCs. Heatmap representation of the main nodes depending of *Fe*_*ext*_ and *O*_2_. The “~” symbol indicates an oscillatory behavior of the variable in the attractor, otherwise the variable value is fixed. The gradient color correspond to the level of activation of the node. In cases where the variable oscillate in a cyclic attractor, the mean of all the values of the node in the cyclic attractor was computed.

### Model predicts growth defects situations

Deficiencies in Fe-S biogenesis are known to cause growth defects. We thus used the model to predict situations in mutants for which *Isc* and *Suf* nodes activities would be turned off, thus affecting growth of bacteria. Out of 54 simple mutants, only the simple *IscR-A* KO (and, more trivially, the *Suf* KO mutant) displayed situations in which nodes *Isc* and *Suf* equaled 0. For combinations of double mutants, only combinations of mutants including *IscR-A* KO or *Suf* KO mutation displayed situtations for which nodes *Isc* and *Suf* were both turned down to 0. The ensemble of these data is available online (https://gitlab.com/Laurent.Tichit/fe-s-cluster-biogenesis-logical-model/tree/master/Results/S4_Fig). For instance, our model predicts that in *Suf* KO there is no Fe-S biogenesis system when *Fe*_*ext*_ =0, in agreement with the fact that *E. coli suf* mutants are sensitive to iron starvation [41] (see Fig 7 A bottom panel). In contrast, while our model predicts a similar deficiency in Fe-S biogenesis in Fe stravation conditions for the *IscR-A* KO mutant, to our knowledge, growth defects of *iscR* mutants had not been reported in the literature. We thus tested if *iscR* mutants displayed growth deficiencies in such conditions by growing *E. coli* WT or *iscR* mutants in medium treated or not with dipyridyl (Dip), a well known iron chelator (Fig 7 B and C). A *suf* mutant strain, deleted for the whole *suf* operon, was also included as a control. Both *iscR* and *suf* mutants grew normally in rich medium (Fig 7 B). As expected, adding 300 *μ*M of Dip iron chelator slightly affected growth of the WT strain as iron is needed for many cellular processes. However, as predicted by the model, this growth defect phenotype was aggravated in the *iscR* and the *suf* mutants, thus validating the model prediction that *iscR* mutants presents growth defects during iron starvation due to its inability to make Fe-S clusters.

**Fig 7.**
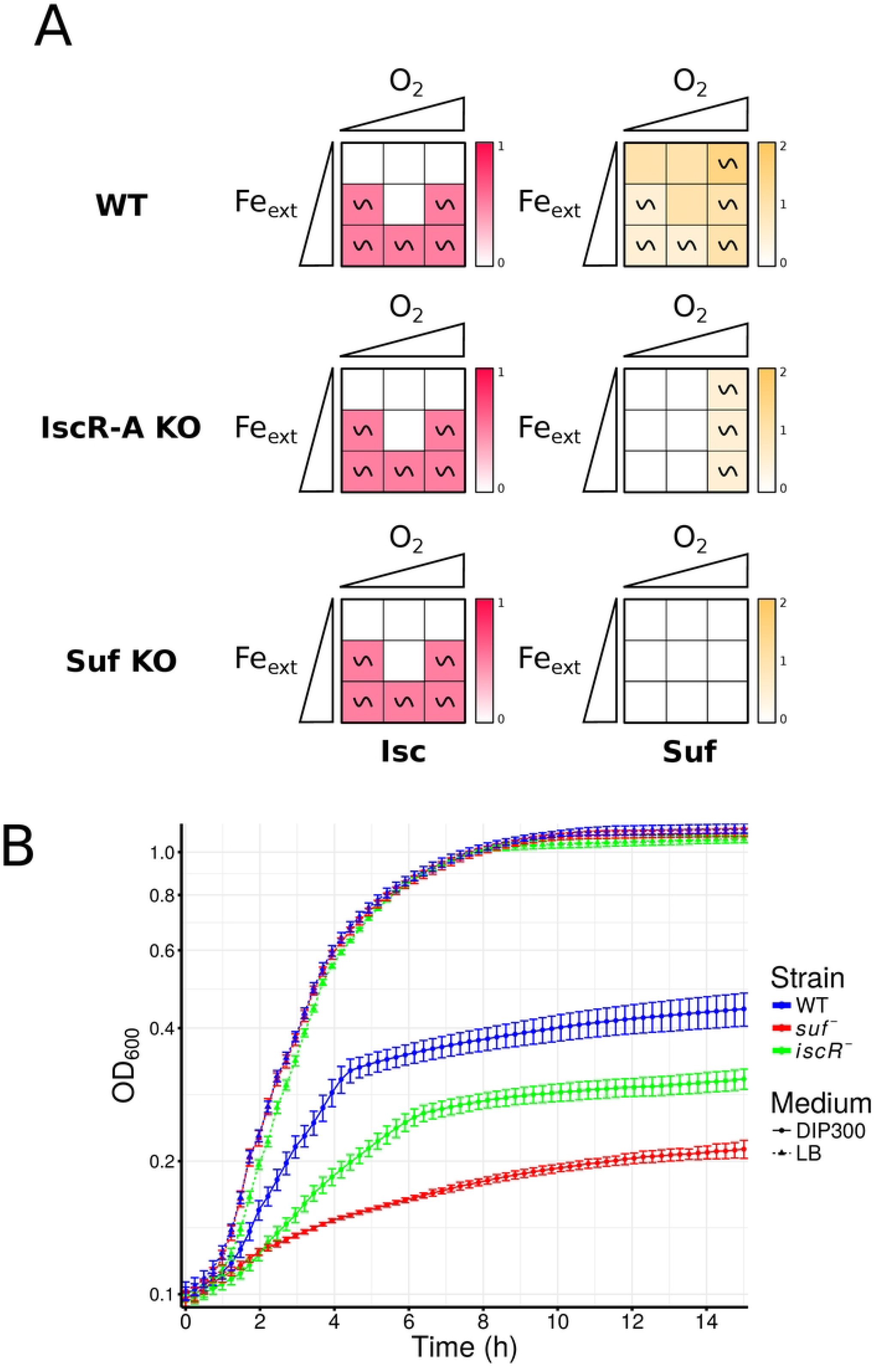
Growth defects predictions and experimental verification. (A) Heatmap representation of situations where neither the Isc and Suf machineries are active in the model. Blue and red circles correspond to the experimentally tested conditions (iron starvation and iron replete conditions respectively). (B) Experimental verification of the predictions. The dashed and filled lines correspond respectively to iron replete and iron deficient conditions. Iron starvation was induced using 300*μ*M of dipyridyl chelator. Growth was measured at OD_600_ for 14h at 37°C using a TECAN microplate reader.

## Conclusion

We have presented here a logical model that enables us to describe for the first time in details how Fe-S biogenesis is finely controlled in response to iron availability and oxidative conditions. It is based on the integration of biological data from literature and genetic experiments that were translated in logical rules and regulatory graph. The simulations of the model recapitulate qualitatively the possible behaviours of the system depending on iron and oxygen availability. We have proposed a visualization of these behaviours that facilitates the detection of changes with the environmental conditions and the different perturbations of the model.

Our model relies on the logical formalism, a discrete mathematical framework that has proved to be particularly successful to understand dynamical features of complex biological processes, and to model and analyze regulatory and signaling networks. Previous models based on ordinary differential equations were built in order to analyse iron homeostasis in *E. coli*, reporting damped oscillations of Fur regulated gene expression [42]. In another modeling driven study, Fe-S cluster biogenesis was predicted to compensate iron import in order to avoid iron toxicity at high iron concentrations [43]. Other models were built in order to analyse oxidative stress in *E. coli*, either for understanding the contributions of AhpCF, KatG and the H2O2 diffusion in the membrane [44], or the effects of carbon deprivation on oxidative stress response [24]. However, the crucial impact of oxidative conditions in coordination with iron concentration in order to mudulate Fe-S biogenesis had not been integrated in the modeling process.

Logical (and more generally discrete) modeling is limited to qualitative results, such as presence or absence of activity of a component in given environmental conditions. Despite this coarse grained abstraction, such a qualitative model captures the most salient properties of the modeled systems. Here it allowed us to summarize the classes of qualitative behaviours depending on environment conditions at the scale of biological processes (modules). We thereby highlight essential features of Fe-S biogenesis. The modular architecture of the network is interesting and could be further exploited.

Indeed, it is believed that the network architecture can help control the network dynamics to some extent. Several methods also attempt to decompose the large network into smaller modules in order to analyze them and then recompose the overall dynamics. This model provides an excellent example for working through these methodological challenges.

Analysis of our model reinforces the classical paradigm of Isc being the homeostatic machinery expressed in favorable conditions in opposition to Suf as the stress responsive machinery (Fig 3). Indeed, perturbing either the oxidative stress or the iron homeostasis responses is predicted to generally induce expresssion of Suf and to turn down Isc (5). However, the analysis of our model highlights interesting complexities.

Our model predicts that expression of the Isc machinery is correlated with iron availability but independent of oxidative conditions (3). Thus, surprisingly, Isc is also predicted to be present during oxidative stress. In fact, activity of Isc has been observed in cells experiencing protracted oxidative stress, validating our model prediction, although Isc was shown to be not functional for maturation of Fe-S targets in such conditions (Jang & Imlay 2010). One explanation to this somewhat paradoxical behavior may be that one or multiple proteins of the Isc machinery serves other functions than Fe-S biogenesis in presence of oxidative stress. IscS in particular also functions as a cysteine desulfurase for tRNA modifications, an activity that may not be compromised in presence of ROS and that cannot be replaced by SufS (Bühning 2017).

Somewhat surprisingly, it should be noted that the situation that gives rise to Suf highest expression in our model is not the one that is predicted to be the more destabilizing for the ensemble of the system (see Fig 2). These results suggest that while Fe starvation diminishes the burden of oxidative stress by hindering Fenton reaction, it worsens Fe-S biogenesis as it becomes even more difficult to synthesize Fe-S clusters in presence of ROS and in iron starved conditions, rising the need for the Suf machinery.

Interestingly, our model also gives new insights on ATC transporter usage in function of environmental conditions. The model predicts that NfuA expression is anti correlated with that of Isc, and becomes higher in stress conditions or by perturbing the iron homeostasis or oxidative response modules (Fig 3 and Fig 6). Conversely, ErpA is predicted to be expressed in almost all conditions in the WT, with a peak at medium iron and oxygen levels, but to be turned down when input modules are perturbed by KO mutations (Fig 3 and Fig 6).Thus, our model highlights the role of NfuA as a stress response carrier while ErpA behaves as an housekeeping component of Fe-S biogenesis, in a parallel situation to Suf and Isc. This model is in agreement with several observations in the literature: in particular, an *nfuA* mutant has been shown to be required during oxidative stress and iron starvation conditions [7] while *erpA* is essential in normal conditions, as would be expected for an important stress or housekeeping component, respectively. However,carriers usage is probably more versatile than that of Fe-S biogenesis machineries. Indeed, both NfuA and ErpA have been shown to be together required for maturation of a subset of Fe-S protein targets and to be able to interact with either the Suf or Isc machineries [40], and this is reflected in our model by the fact that both carriers are expressed together in all conditions in the WT.

Finally, in the past few years Fe-S biogenesis machineries usage has been shown to have important consequences on the sensitivity to antibiotics, in particular aminoglycosides [45]. Indeed, such antibiotics enter the cell through the proton motive force generated by respiratory complexes I and II (Nuo and Sdh). As respiratory complexes are composed of numerous iron-sulfur containing proteins, their activity is heavily dependent on Fe-S biogenesis in the cell, and in particular only Isc activity enables sufficient respiratory complexes maturation to allow aminoglycosides entry. As a consequence, iron starvation, in which Suf is used instead of Isc, induces resistance to aminoglycosides, and RyhB has a major role in promoting this resistance phenotype through its inhibition of Isc synthesis [46]. By enabling to precisely predict machineries usage in function of the environmental conditions, our model should also predict situations for which the bacterium becomes resistant to aminoglycosides. For instance, the Fur mutant is predicted to be resistant to aminoglycosides regardless of the growth conditions, as Isc is down-regulated (see Fig 5), a phenotype that has been validated in laboratory growth conditions [45].

## Materials and methods

### Logical modeling

We chose to work within the logical framework which offers a qualitative description of the molecular mechanisms, and is therefore well suited to elucidate main processes of the cell functioning and to pinpoint key biological components controlling the modelled process. It relies on a **Regulatory Graph** (RG), a signed labelled directed graph whose nodes represent the biochemical species, and directed signed edges linking pairs of nodes represent the regulations (activations or inhibitions). The selection of molecular players to be included in the RG and their mutual interactions are based on an extensive analysis of the literature and on our current understanding of the molecular mechanisms controlling Fe-S biogenesis (schematized in the biological map Fig. S1). To complete this model description and obtain a dynamical and predictive model, we attached to each node a discrete variable representing its activity level that reflects the node’s ability to regulate its targets in the network. With a Boolean variable, the component is active (ON/1) or inactive (OFF/0). In some situations, more than two levels are necessary to distinguish different regulatory roles of the component in the network and the variable is multilevel. The threshold at which a regulation takes place is indicated as a label on the edge (1 by default). Vectors encompassing values for all components are called logical states. The impacts of combinations of regulators on the state of a component are encoded through logical rules (cf [47]) expressed with logical operators AND, OR, NOT.

Given a logical state, the logical rules determine which components are able to change their value. We use an asynchronous dynamics: only one node can update at a time. Hence, two successive states differ by only one component, and a state has as many successors as the number of components called to update. This dynamics is thus non deterministic, and all the possible asynchronous trajectories can be represented in the **State Transition Graph** (STG), a directed graph whose nodes are the logical states of the network, and edges connect successive states.

To characterize the long-time behaviours of the system, we identify the attractors of the model which are the terminal strongly connected components (SCC) of the STG, i.e. the sets of states in which the dynamics is trapped. Their gene expression patterns reveal the activities of components at these asymptotic regime. We distinguish two types of attractors, the stable states constituted of a unique node (all the components are stable); and the complex (or cyclical) attractors containing more than 2 nodes (some components display oscillations). The identification of complex attractors is often computationally difficult because of the combinatorial explosion of the STG. So we take advantage of functionalities implemented in GINsim, in particular we generate the **Hierarchical Transition Graph** (HTG) that is a compaction of the STG [48].

### Logical modeling Software GINsim

We used the freely available software GINsim (Gene Interaction Network simulation, http://ginsim.org) for the edition, analysis and simulation of the regulatory graph [49]. Once the model defined, several simulation tools are proposed to simulate the State Transition Graph, the Hierarchical Transition Graph, do reduction of model or compute the stable states. GINsim also enables the simulation of mutants (loss-of-function, ectopic gene expression, and combinations) by blocking the levels of expression of the corresponding variables. To further ease the analysis of multiple perturbations, we have written a set of scripts in Python and R, which iteratively compute the behaviour of our model for every input values and mutants considered, and generate heatmaps representing their behaviour.

### Growth assays

Sensitivity to iron starvation was measured as follows: overnight stationary phase culture grown in LB media were diluted 1:500 in 3 mL LB without or with 300 *μ*M DIP. Cells were grown with shaking at 37°C for 9 hours, before being diluted 1:100 in a 96 well plate containing either LB with or without 300 *μ*M DIP. Growth of the different strains were measured by reading the OD600 of each well during 14 h at 37 °C on a TECAN Spark 510M.

### Reproducibility of the results

In order to allow reproducing the results, we have developed an application allowing to:

1. load the logical model (in the GINsim format) corresponding to the current study
2. after choosing N nodes of interest, build a set of mutant models (which contains the “wild type” and all the single mutants, double mutants, …, N-uple mutants)
3. compute the dynamics of each model using GINsim
4. choose the readouts (the variables of interest representing either the expression level, activity or discrete concentrations) to visualize, and optionally search for mutants which may lead to the desired phenotype (e.g. by default, low levels of Isc activity and Suf expression in the attractor)
5. finally, generate the heatmap representation of the asymptotic behaviors of the readouts for each mutant.

This application is installable directly from source from gitlab and can be run via executing a docker file containing all the required dependencies. It is available at https://gitlab.com/Laurent.Tichit/fe-s-cluster-biogenesis-logical-model/.

## Supporting information

**S1 Text**. Full model description.

**S1 Table**. Description of the variables

**S2 Table**. Description of the attractors of the model

**S3 Table**. Model validation

**S2 Text**. Model prediction.

## Notes

### Competing Interest Statement

The authors have declared no competing interest.

